# Detecting Insulitis in Type 1 Diabetes with Ultrasound Phase-change Contrast Agents

**DOI:** 10.1101/2020.10.28.359687

**Authors:** David G. Ramirez, Awaneesh K. Upadhyay, Vinh T. Pham, Mark Ciccaglione, Mark A Borden, Richard K.P. Benninger

## Abstract

Type 1 diabetes (T1D) results from immune infiltration and destruction of insulin-producing β-cells within the pancreatic islets of Langerhans (insulitis), resulting in loss of glucose homeostasis. Early diagnosis during pre-symptomatic T1D would allow for therapeutic intervention prior to substantial loss of β-cell mass at T1D onset. There are limited methods to track the progression of insulitis and β-cell mass decline in pre-symptomatic T1D. During insulitis, the islet microvasculature increases permeability, such that sub-micron sized particles can extravasate and accumulate within the islet microenvironment. Ultrasound is a widely deployable and cost-effective clinical imaging modality. However, conventional microbubble contrast agents are restricted to the vasculature. Sub-micron sized nanodroplet (ND) phase-change agents can be vaporized into micron-sized bubbles; serving as a circulating microbubble precursor. We tested if NDs extravasate into the immune-infiltrated islet microenvironment. We performed ultrasound contrast-imaging following ND infusion in NOD mice and NOD;Rag1ko controls, and tracked diabetes development. We measured the biodistribution of fluorescently labeled NDs, with histological analysis of insulitis. Ultrasound contrast signal was elevated in the pancreas of 10w NOD mice following ND infusion and vaporization, but was absent in both the non-infiltrated kidney of NOD mice and pancreas of Rag1ko controls. High contrast elevation also correlated with rapid diabetes onset. In pancreata of NOD mice, infiltrated islets and nearby exocrine tissue were selectively labeled with fluorescent NDs. Thus, contrast ultrasound imaging with ND phase-change agents can detect insulitis prior to diabetes onset. This will be important for monitoring disease progression to guide and assess preventative therapeutic interventions for T1D.

**Significance:** There is a need for imaging methods to detect type1 diabetes (T1D) progression prior to clinical diagnosis. T1D is a chronic disease that results from autoreactive T cells infiltrating the islet of Langerhans and destroying insulin-producing β-cells. Overt disease takes years to present and is only diagnosed after significant β-cells loss. As such, the possibility of therapeutic intervention to preserve β-cell mass is hampered by an inability to follow pre-symptomatic T1D progression. There are immunotherapies that can delay T1D development. However identifying ‘at risk’ individuals, and tracking whether therapeutic interventions are impacting disease progression, prior to T1D onset, is lacking. A method to detect insulitis and β-cell mass decline would present an opportunity to guide therapeutic treatments to prevent T1D.

## Introduction

Type 1 diabetes (T1D) is caused by infiltration of auto-reactive immune-cell into the islets of Langerhans in the pancreas (insulitis) and destruction of insulin-secreting β-cells. The subsequent loss of glucose homeostasis requires lifelong insulin therapy, with T1D subjects still at elevated risk of chronic diabetes complications and hypoglycemia-induced coma or death. Prior to clinical onset, there exists an asymptomatic phase of many years where autoimmunity and insulitis progresses but glucose homeostasis is maintained (pre-symptomatic T1D) (1). However, patients with T1D are not diagnosed until presentation of hyperglycemia, where the majority of β-cell mass (>80%) has been lost. As such this pre-symptomatic phase of T1D, where β-cell mass remains, provides an ideal window for the therapeutic prevention of T1D (2).

Preservation of β-cell mass would delay the onset of T1D and reduce the incidence of diabetic complications and hypoglycemia after T1D onset. There has been some limited success in trials applying immunomodulatory agents designed to preserve β-cell mass and reverse or prevent T1D (3). For example, antiCD3 has recently been applied in first-degree relatives at high risk for T1D and showed a significant delay and reduced incidence of T1D onset (4). However prevention was not complete and these subjects were still significantly advanced in T1D progression, showing signatures of dysglycemia. Application of various therapies at T1D onset similarly show only a temporary preservation of β-cell mass (as measured by c-peptide release), with a broad heterogeneity in treatment response (4). As such earlier therapeutic treatment is desirable.

There are currently very limited means to determine whether a subject shows pre-symptomatic T1D and thus will develop T1D. Furthermore, there are no means to assess the success or failure of therapeutic interventions beyond the onset of T1D itself. The presence of multiple islet-associated autoantibodies in circulation can determine risk of developing T1D within a multi-year time frame (5, 6). However, islet-associated autoantibodies are not pathogenic and cannot inform on successful therapeutic treatments. As such there is a need to develop new methods to detect the progression if insulitis and β-cell mass decline during the pre-symptomatic phase of T1D. Furthermore, at clinical onset there is a high incidence of diabetic ketoacidosis (DKA) that can have severe health consequences (7). Thus identifying ‘at risk’ subjects and monitoring their underlying disease progression will also be important to mitigate the consequences of DKA.

Several imaging approaches have been explored to detect insulitis and β-cell mass decline (8–11). During insulitis, the islet microvasculature increases permeability (11, 12). As such, the enhanced permeability and retention (EPR) effect can be used, where sub-micron sized agents can accumulate in the inflamed islet microenvironment. This effect has been demonstrated using MRI contrast agents, in both mouse models of T1D and human T1D (13, 14). While highly promising, MRI modalities do carry some limitations in terms of deployability and cost effectiveness. Ultrasound imaging modalities can be widely deployed, are costeffective and provide real-time imaging. Contrast-enhanced ultrasound uses gas filled microbubbles and non-linear signal detection. However conventional microbubble contrast agents are restricted to the vasculature. Sub-micron sized nanobubble ultrasound contrast agents have been developed. These agents act via the EPR effect to accumulate in tumors and inflamed tissue, including the immune infiltrated pancreas (15, 16). However one potential limitation with sub-micron sized bubbles is their low scattering cross section, requiring a larger number of bubbles to be infused to generate measurable signal.

Nanodroplet phase-change agents are a class of sub-micron sized ultrasound contrast agents (17, 18). Nanodroplets consist of a super-heated gas core and folded lipid shell that are stable at body temperature. However, within an acoustic beam these nanodroplets can vaporize into micron-sized bubbles that provides significant ultrasound contrast. As such these nanodroplets can act as a circulating nanoscale microbubble ‘precursor’ that can be ‘activated’ by an acoustic beam. Nanodroplets can be readily formed by condensing conventional microbubble agents (19), including clinically approved agents (20). Given their sub-micron size, nanodroplets have been demonstrated to follow the EPR effect and accumulate in tumors and generate measurable ultrasound contrast signal (21, 22). However, whether nanodroplets can similarly accumulate in inflamed tissues and whether this accumulation can be measured and used for disease monitoring has been lacking.

In this study, we tested whether nanodroplet phase change contrast agents would specifically accumulate within inflamed islets of Langerhans in mouse models of T1D; and the degree to which this accumulation depends on the level of insulitis. We further tested whether this accumulation could be measured non-invasively via ultrasound contrast imaging, as an increase in contrast signal following nanodroplet vaporization, and how this correlated with diabetes onset.

## Results

### Sub-micron nanodroplets accumulate and can be vaporized within the pancreas of NOD mice

Sub-micron sized particles selectively accumulate in the islets of Langerhans in mouse models of type1 diabetes (T1D) via the enhanced permeability and retention (EPR) effect that can occur in inflamed tissues (12, 15). Nanodroplet phase change agents are sub-micron sized that can accumulate in tumors, and can be vaporized within the tissue to provide significant ultrasound contrast. We first tested whether nanodroplets would similarly accumulate within immune-infiltrated islets of Langerhans and provide significant ultrasound contrast following vaporization. Size-isolated microbubbles were prepared and condensed under high pressure and low temperature into sub-micron-sized droplets (Fig. 1a). We utilized 10w female non-obese diabetic (NOD) mice where the immune-infiltrated pancreas and non-immune infiltrated kidney can be readily identified under B-mode imaging, with the spleen and kidney serving as landmark for pancreas identification (Fig. 1b).

**Figure 1.**
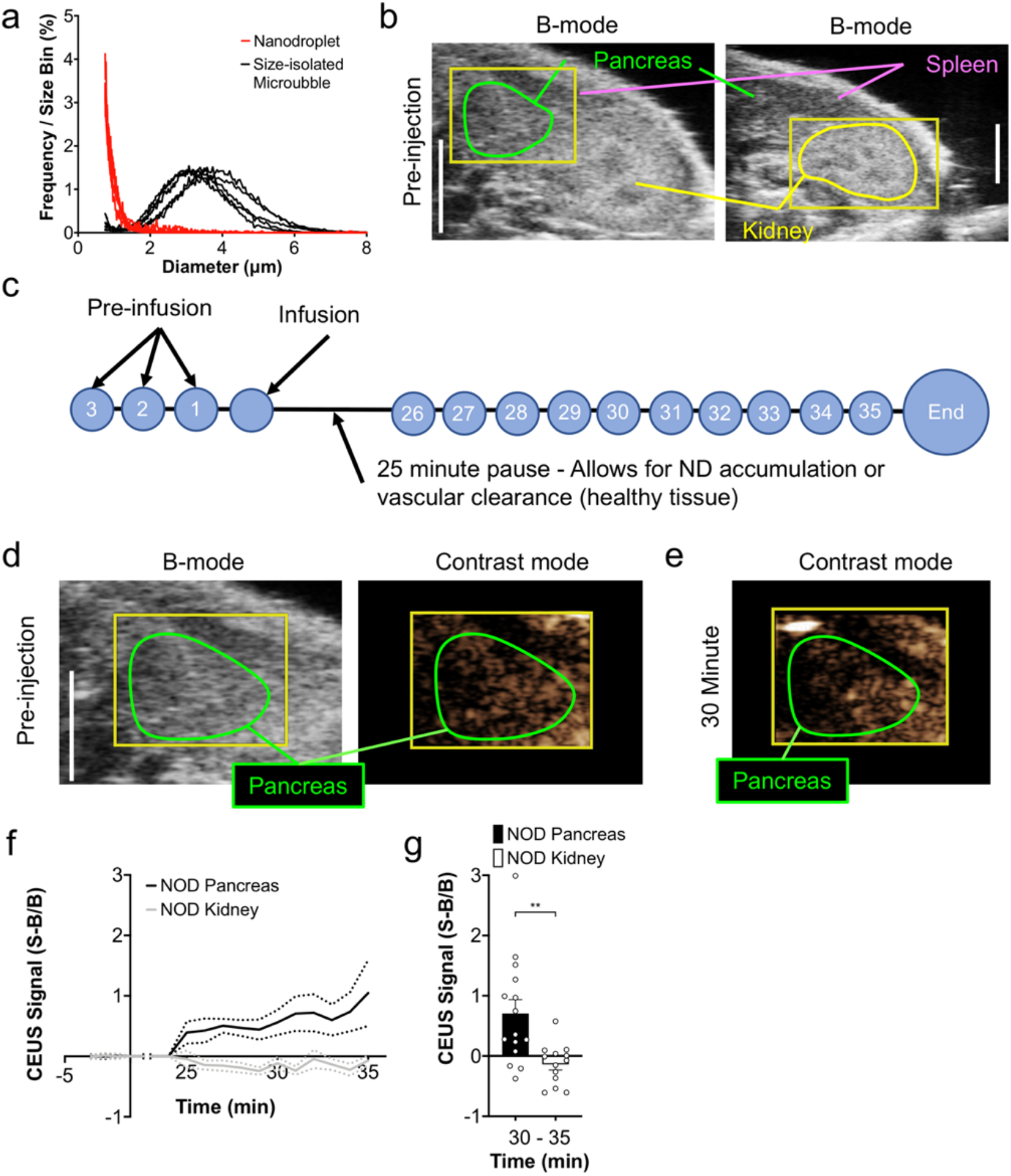
Nanodroplet infusion and vaporization increases contrast in the pancreas prior to T1D. B) Size distribution of size-isolated microbubbles prior to condensation and of nanodroplets following condensation. (B) Representative B-mode images of the abdomen, indicating the pancreas (green) and kidney (yellow), as well as the spleen anatomical landmarks used to identify the pancreas. (C) Schematic of nanodroplet imaging protocol, where time courses are recorded prior to nanodroplet infusion (pre-infusion). After nanodroplet infusion, acquisition is paused to allow the nanodroplets to accumulate inn tissues with permeable microvasculature and clear from the bloodstream. Following the pause, time courses are then recorded for ten minutes during which droplets are vaporized. (D) Representative B-mode and sub-harmonic contrast images of pancreas (green) before nanodroplet infusion. (E) Representative sub-harmonic contrast ultrasound image of pancreas (green) 28 minutes after nanodroplet infusion. (F) Time-course of mean contrast signal, normalized to background, in the pancreas and kidney of 10-week-old female NOD mice following nanodroplet infusion and vaporization. (G) Mean contrast signal averaged between 30-35 minutes following nanodroplet infusion. Dashed lines in F represent 95% CI, error bars in G represent s.e.m. Data in F, G representative of n=15 mice (pancreas) and n=13 mice (kidney). The p-value in G is 0.0034 (**p<0.01), comparing groups indicated (unpaired Student’s t-test).

To measure whether nanodroplets accumulate within the islet of NOD mice, we designed a protocol to separate nanodroplets in circulation from tissue accumulation (Fig. 1c). First, a defined region covering either the pancreas or kidney was chosen (Fig. 1a). Next, a series of sub-harmonic contrast mode images were acquired for 3 minutes prior to nanodroplet infusion, forming the background time courses. Following nanodroplet infusion acquisition of B-mode or contrast mode images was paused for 25 minutes to allow maximal accumulation within inflamed tissues. This pause is based upon both size-isolated microbubbles and nanobubble contrast agents clearing from circulation within 5-10 minutes (11). Following this pause, a series of sub-harmonic contrast mode images were acquired for 10 minutes while the defined region was insonified and nanodroplets vaporized. We compared the contrast signal within the pancreas or kidney prior to infusion (Fig.1d) with the contrast signal during nanodroplet vaporization (Fig. 1e). Following nanodroplet infusion, increased contrast signal over the background was observed in the pancreas within ~3 minute of insonifying the pancreas region (Fig. 1e, f). After this time there was no further significant elevation in contrast signal. Within the kidney we observed no such elevation in contrast signal while insonifying the kidney region (Fig. 1f). The increase in contrast signal within the pancreas 5 minutes after insonficaiton was significantly elevated compared to the contrast signal within the kidney (Fig. 1g). These results suggest that nanodroplets are specifically accumulating within the pancreas where immune-infiltrated islets of Langerhans are located, and these nanodroplets can be vaporized within the tissue to provide measurable ultrasound contrast.

### Sub-micron nanodroplet accumulation is specific to animals that develop diabetes

Given lack of contrast elevation in the kidney of NOD mice, we next asked whether nanodroplet accumulation was specifically associated with immune cell infiltration. We compared 10w female NOD mice to age-matched and sex-matched immune deficient NOD;Rag1ko (Rag1ko) mice. In each case we followed the same imaging protocol as before, focused on the pancreas (Fig. 2a, 2b). Rag1ko mice do not develop mature T and B cells and thus do not show any immune cell infiltration into the islet and do not develop diabetes. As above, in NOD mice we observed a substantial elevation in contrast signal within ~3 minutes of insonification, indicating vaporization of accumulated nanodroplets in the pancreas (Fig.2a,c). However, in the pancreas of NOD Rag1ko mice we did not observe any contrast elevation following insonification of the pancreas (Fig. 2b,c), indicating lack of any nanodroplet accumulation and vaporization. The increase in contrast signal within the pancreas 5 minutes after insonficaiton was significantly elevated in NOD mice compared to in Rag1ko mice (Fig. 2d).

**Figure 2.**
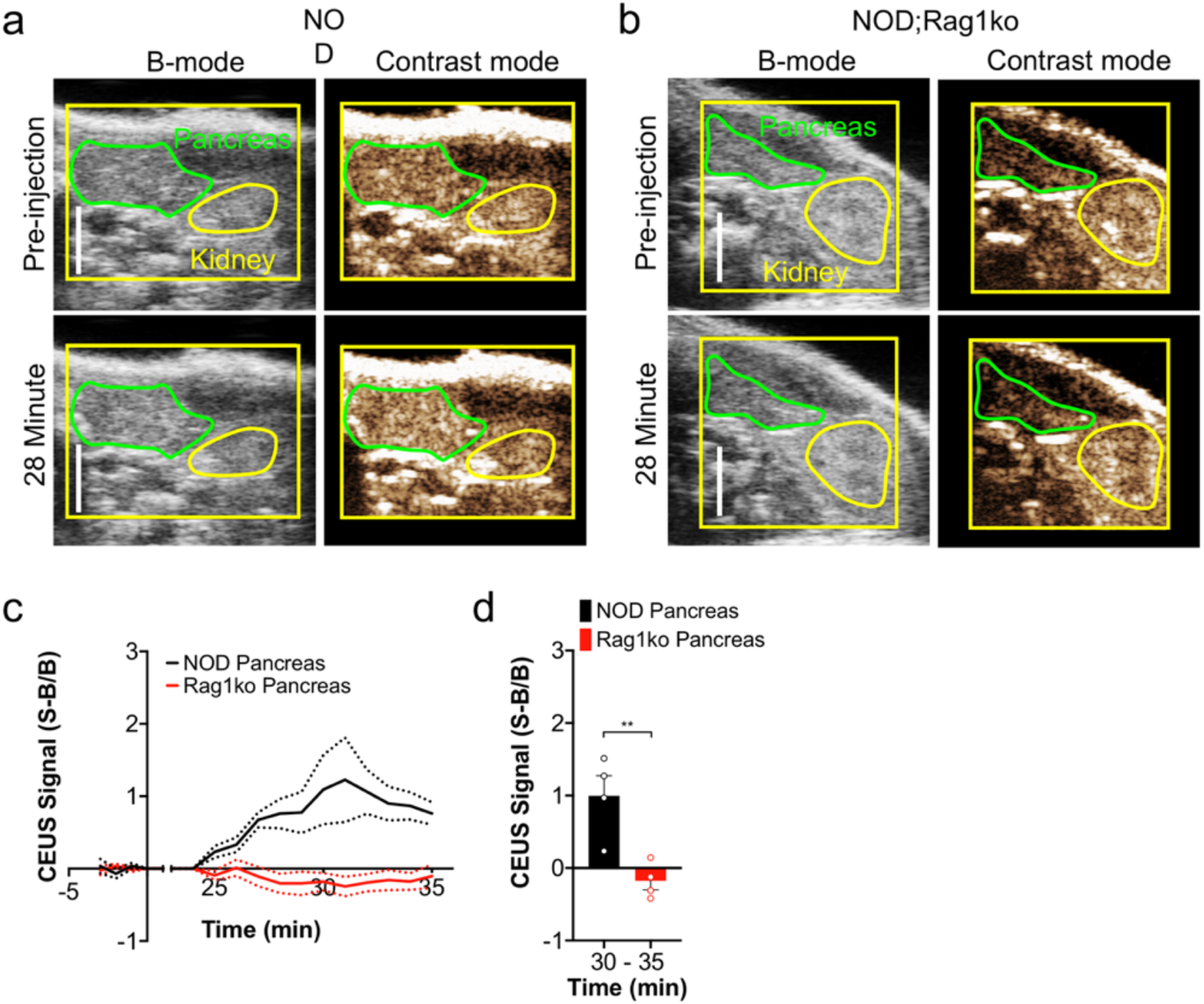
Nanodroplet infusion and vaporization increases contrast dependent on insulitis and T1D. (A) Representative B-mode and sub-harmonic contrast images of the pancreas (green) and kidney (yellow) before and after nanodroplet infusion in 10-week-old female NOD mice. (B) As in A, but for 10 week old immune-deficient female NOD;Rag1ko (Rag1ko) mice. (C) Time-course of mean contrast signal, normalized to background, in the pancreas of 10-week-old female NOD mice versus 10-week-old female Ragk1o mice following nanodroplet infusion and vaporization. (D) Mean contrast signal averaged between 30-35 minutes following nanodroplet infusion. Dashed lines in C represent 95% CI, error bars in D represent s.e.m. Data in C, D representative of n=4 mice (NOD) and n=4 mice (Rag1ko). The p-value in D is 0.0083 (**p<0.01) comparing groups indicated (unpaired Student’s t-test).

While NOD mice show significant insulitis at age 10 weeks of age (23) they do not generally become hyperglycemic until after 12 weeks of age. Furthermore there is a wide range of ages at which diabetes develops, from 12 weeks to up to 40 weeks of age. Glucose measurements of female NOD mice are consistent with these reported numbers, where mice developed diabetes between 12 weeks and >36 weeks of age (Fig. 3a). Interestingly, we observed a significant correlation between the mean elevated contrast signal in the pancreas upon nanodroplet vaporization with the time from the measurement at which diabetes arises (Fig. 3b). Thus, mice that show elevated ultrasound contrast, associated with increased nanodroplet accumulation within the pancreas, progress more rapidly to diabetes.

**Figure 3.**
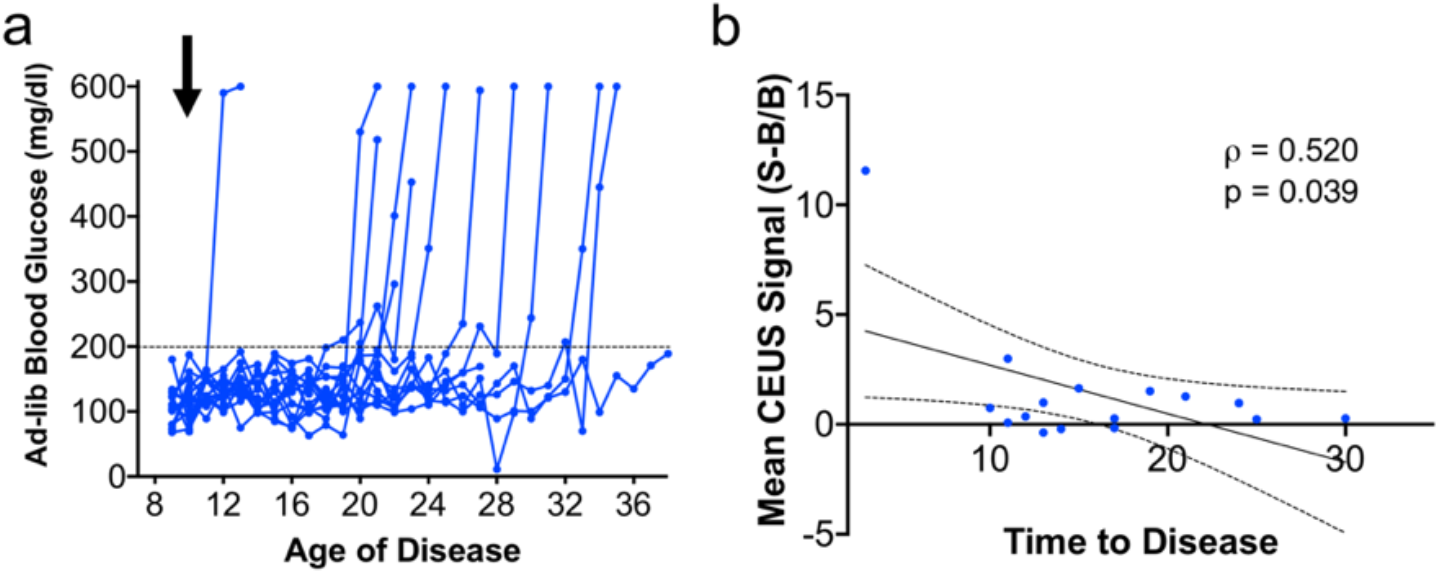
Nanodroplet contrast elevation correlates with T1D onset in NOD mice. (A) Time course of ad-lib measured blood glucose in the NOD mice that received nanodroplet infusion. Blue arrow indicates time points of nanodroplet infusion and ultrasound scans. (B) Scatterplot of mean contrast elevation against the time to diabetes onset, for NOD mice. Data in A, B represents n=16 NOD mice. A mixed-effects model was used to assess the statistical significance and generate the regression in B.

### Nanodroplet accumulation within pancreas of NOD mice is restricted to the islet environment

The pancreas contains 2 distinct compartments: the islets of Langerhans (endocrine pancreas) where immune cell infiltration occurs in NOD mice, and the exocrine pancreas where immune cell infiltration is minimal. To determine the biodistribution of nanodroplets within the pancreas undergoing insulitis, we infused DiO-labeled polydisperse nanodroplets into 10w NOD mice, and vaporized these nanodroplets in a subset of mice following the same procedure as above. Following nanodroplet vaporization (or a pause for those where nanodroplets were not vaporized) we dissected and sectioned the pancreas for histological analysis. Following infusion and vaporization, nanodroplets generated from (unlabeled) polydisperse microbubbles show elevated contrast signal in the pancreas, but not in the kidney (Fig. S1), similar to that measured in nanodroplets generated from size-isolated microbubbles.

We first assessed the coverage of DiO fluorescence in pancreas; within the islets, in regions close to the islet (within 200μm, ‘near islet’) and within the exocrine pancreas far (>500μm) from the islet (Fig. 4a-c). We also compared the coverage of DiO fluorescence in these pancreas regions for vaporized and non-vaporized nanodroplets (Fig. 4a,d). In 10w NOD mice with vaporized NDs, both the islet and regions close to the islet within the pancreas showed significantly more fluorescent coverage compared to exocrine pancreas far from the islet (Fig. 4e). We did not observe any difference in fluorescence coverage between islet and ‘near islet’ regions (Fig. 4e). The fluorescence coverage within islet regions was marginally greater (p=0.06) in the pancreata of animals were nanodroplets were vaporized, compared to animals were there was no vaporization (Fig. 4f). Similarly, the fluorescence coverage within ‘near islet’ regions was significantly greater in the pancreata of animals were nanodroplets were vaporized, compared to animals were there was no vaporization (Fig. 4g). Thus, nanodroplets target the islet and near islet regions more favorably compared to exocrine regions in NOD mice.

**Figure 4.**
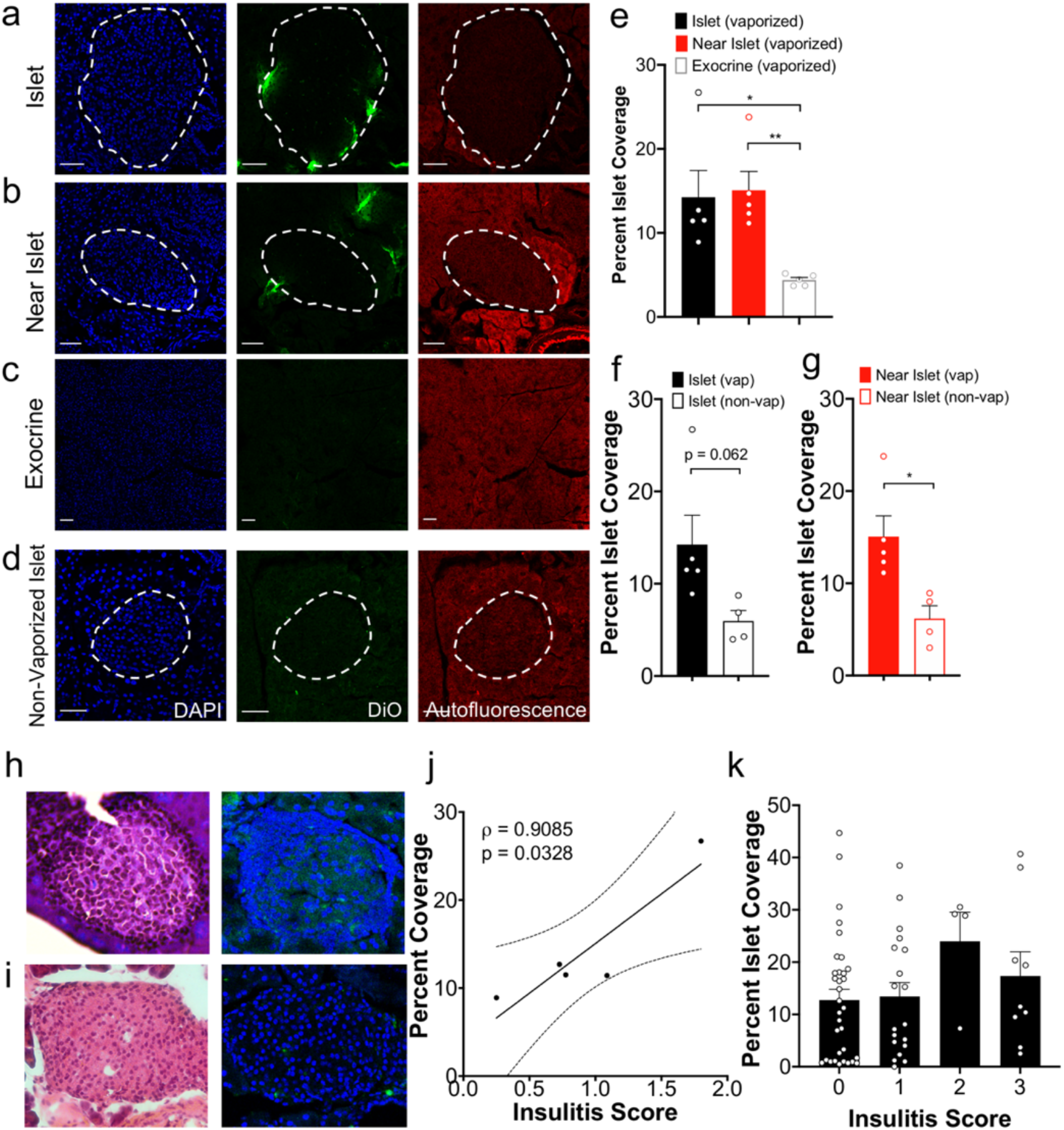
Histological assessment of Nanodroplet accumulation within the islets in T1D. (A) Representative confocal images of an islet within a pancreas section of 10-week-old female NOD mouse following DiO-labeled nanodroplet infusion (green). Islet is circled with a dotted line, as determined from autofluorescence (red) and DAPI-labeling morphology (blue). (B) as in A for “Near Islet” (<200 μm from the islet) regions within the pancreas. (C) As in A for exocrine region far (>500 μm from the islet). (D) as in A but for female 10-week-old NOD mice that did not receive ultrasound scans and therefore had no nanodroplet vaporization. (E) Mean DiO coverage in the islet, near islet regions and exocrine tissue in 10-week-old female NOD mice where nanodroplets were vaporized. (F) Mean DiO coverage in the islet (from 10-week-old female NOD mice in which nanodroplets were vaporized or non-vaporized. (G) As in F in near islet regions. (H) Representative image of hematoxylin and eosin (H&E) stained pancreas sections of 10w NOD mice, together with DiO fluorescence in neighboring tissue section. Islet circled with a dashed line. (I) Scatterplot of the mean DiO fluorescent coverage of islets within the pancreas vs the mean insulitis score within the pancreas, for NOD mice. (J) Mean DiO fluorescent coverage of islets within the pancreas that show insulitis scores of 0, 1, 2, or 3 (see methods). Error bars in E, F, G, J represent s.e.m. Trend line in I indicates linear regression with 95% confidence intervals. Data in E represents n=5 mice (89 islets and 25 exocrine regions). Data in F,G (Vaporized) represents n=5 mice (89 islets) and (Non-Vaporized) represents n=4 mice (37 islets). Data in I represents n=5 NOD mice (73 islets). Data in J represents n=5 NOD mice (65 islets). Scale bar represents 50 μm in A-D. *p<0.05, **p<0.01, comparing groups indicated (Unpaired Student’s t-test for data in E, F, G; ANOVA for data in J). A mixed-effects model was used to assess the statistical significance and generate the regression in I.

To determine if nanodroplet accumulation is dependent on the level of insulitis, and thus underlying disease, we scored the level of insulitis in H&E-stained sections adjacent to those in which we measured DiO fluorescent coverage (Fig. 4h). In pancreata from 10w NOD mice the mean DiO coverage across islet regions of the pancreas was significantly correlated with the mean insulitis score across islets of the same pancreas (Fig. 4i). Because we used adjacent sections when performing histology, we then examined insulitis on an islet-by-islet basis. Interestingly, there was no relationship between islet insulitis and fluorescent coverage (Fig. 4j): islets in which no insulitis was observed (score 0) showed similar DiO fluorescent coverage as islets in which substantial insulitis is observed (score 3). This is consistent with other reports in which the vascular permeability reflects the overall diseased state of the pancreas rather that the local level of disease (15). Consistent with measurements above, there was no relationship between fluorescent coverage and insulitis for non-vaporized nanodroplets, whether analyzed by pancreas or by islet (Fig.S2). Taken together, this demonstrates that within the NOD mouse nanodroplets specifically localize to the islet regions of the pancreas dependent on the overall immune-infiltrated state of the pancreas.

## Discussion

There are limited means for detecting insulitis and β-cell mass decline during the pre-symptomatic phase of type1 diabetes (T1D). A means to detect disease progression during this phase would guide therapeutic treatment during a window in which most β-cell mass remains, together with assessing the success of therapeutic intervention on disease progression prior to diabetes onset. The islet micro-vasculature increases permeability as a result of immune cell infiltration (insulitis). Nanodroplets are submicron in sized and thus can access permeable tissue (22), and following vaporization into microbubbles show a high scattering cross-section. Our goal was to test if sub-micron sized nanodroplets (NDs) would specifically accumulate within immune-infiltrated islets during this pre-symptomatic phase; and whether these nanodroplets could be vaporized to provide elevated ultrasound contrast specifically in the pancreas (Fig. 5).

**Figure 5.**
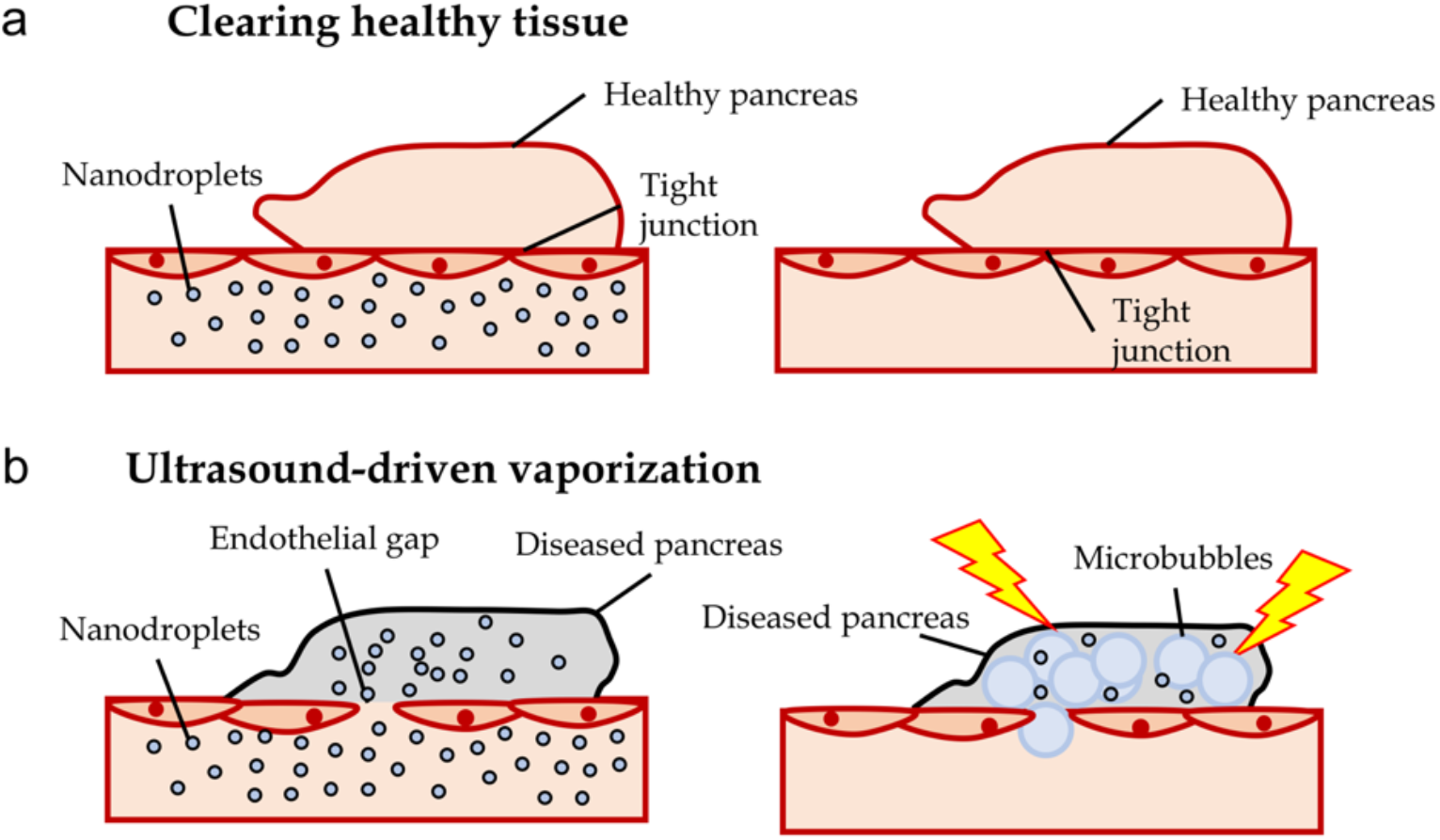
Summary of experiments demonstrating nanodroplets accumulate in immune-infiltrated islets and show increase contrast signal within the pancreas after vaporization. (A) After nanodroplets are infused the healthy islet microvasculature is not permeable (left), and nanodroplets are cleared from the blood stream (right). (B) Immune-infiltrated islets in pre-symptomatic T1D have high vascular permeability and infused nanodroplets accumulate in the tissue (left). Following clearing from the vasculature, vaporization of accumulated nanodroplets, provides a source of increased sub-harmonic contrast (right). This contrast signal therefore provides a measure of insulitis and thus pre-symptomatic T1D.

### Nanodroplet accumulation in pre-symptomatic T1D correlates with insulitis and diabetes progression

Our results demonstrated that nanodroplets specifically accumulate the pancreatic islets during the progression of insulitis in pre-symptomatic T1D. We observed significant contrast elevation within the pancreas following infusion and clearing from circulation indicating that droplets had accumulated and vaporized within the pancreas. Notably a contrast elevation was absent in non-infiltrated organs(e.g. kidney) within the NOD model of T1D (Fig.1). Contrast elevation was also absent in the pancreas of immune-deficient Rag1ko mice that lack insulitis and do not develop diabetes (Fig.2). Thus we conclude that nanodroplets specifically accumulate within the organs in which immune infiltration and inflammation occur. Via histological analysis we further determined that within the pancreas accumulation was specific to the regions that underwent insulitis. The correlation between nanodroplet accumulation within islet regions with the level of insulitis across the pancreas further supports the specificity of nanodroplet accumulation.

While we observed a strong correlation between insulitis and nanodroplet accumulation on a pancreas wide basis, we did not see any link between insulitis and accumulation on an islet wide basis. That is within a pancreas that on average showed high levels of insulitis, islets with lower or even absent insulitis still showed significant nanodroplet accumulation (Fig. 4). When examined, this has been observed with other contrast agents that extravasate and accumulate within the islet in models of T1D (13, 15, 24). Thus while islet microvascular permeability is a good indication of pre-symptomatic T1D (13, 14), it does not necessarily reflect spatial variations in insulitis across the pancreas, at least in mouse models of T1D. Testing whether there is a link between spatial variations in insulitis and islet microvascular permeability in human T1D is still needed. Nevertheless, the specific mechanisms leading to increased islet permeability concurrent with insulitis is still unknown, and will be needed to identify the mechanisms underlying T1D that are reflected by such contrast accumulation measurements, such as the method described here.

Thus nanodroplet accumulation and contrast elevation upon vaporization provides a non-invasive method to assess the state of insulitis across the pancreas in T1D, prior to T1D onset at a point where animals are euglycemic.

### Nanodroplet accumulation predicts disease effectively

A key goal for imaging diagnostics for T1D is to detect signatures of pre-symptomatic T1D early, where significant beta cell mass remains and thus therapeutic treatment can be most effective (2). Predicting T1D onset may also avoid complications associated with diabetic ketoacidosis (7). We detected signatures of insulitis prior to diabetes onset, where animals were euglycemic, that were absent in immunodeficient mice that did not develop diabetes. More importantly the contrasts signal we measured following nanodroplet vaporization also correlated with the time to diabetes onset (Fig. 3): those animals that showed high contrast elevation proceeded to diabetes more rapidly. This indicates a further ability to predict the progression to T1D, which will be important to determine the time for therapeutic intervention and to assess the success of this intervention. Current predictions using circulating autoantibody measurements can only predict diabetes onset within a 10-year window (5). Future work will need to test whether nanodroplet accumulation, vaporization and contrast elevation can longitudinally track pre-symptomatic T1D, and include earlier time points, to determine whether the success of therapeutic treatments can be predicted. Furthermore future work will need to determine whether c-peptide preservation and T1D prevention is more effective at the stages at which signatures of pre-symptomatic T1D are detected using the method described here, compared to e.g. presence of dysglycemia or following T1D onset. Such rigorous analysis has not been performed using any contrast agent.

Thus nanodroplet accumulation and contrast elevation provides a method to predict the rapid onset of T1D, which may be useful for therapeutic treatment.

### Relevance to clinical application

While we have demonstrated high utility for nanodroplet-based ultrasound measurements in mouse models of T1D there are important considerations to be made for clinical translation to human pre-symptomatic T1D detection and tracking. Iron-oxide nanoparticle MRI contrast agents selectively accumulate within the pancreata of animal models of T1D, as well as recent onset human T1D (14, 25). Thus in human T1D there is increased microvascular permeability at diabetes onset, although prior to onset the time-course of islet microvascular permeability is unknown. This compared to increased microvascular permeability in the NOD mouse as early as 4-6 weeks (15). Furthermore, while the EPR effect occurs in human T1D, it will be important to know how permeability compares to insulitis in the human pancreas.

Microbubble contrast agents are clinically approved, including within pediatric subjects (26). While use for pancreas indications is not approved several off-label studies have been reported for pancreatic cancer and pancreatitis (27, 28). Thus imaging ultrasound contrast in the pancreas in human subjects is feasible. While we utilized a small animal ultrasound machine in this study, nanodroplets can be vaporized using clinical ultrasound frequencies (Alec 2020, Dayton 2016(?)). Further, higher frequencies may be more suitable for pediatric subjects and the pancreas can be imaged using higher frequencies via endoscopic ultrasound (28).

While microbubble formulations such as DEFINITY^®^ are clinically approved nanodroplets have not been to date. However, nanodroplets can be readily generated from microbubble contrast agents with minimal intervention (19): here we condensed microbubble using low temperature and high pressure only. Further we demonstrated nanodroplets generated from a DEFINITY-like polydisperse formulation of microbubbles accumulated specifically within the islets of the pancreas and generated ultrasound contrast upon vaporization (Fig. 4, Fig. S1). Nanodroplets have also been generated from DEFINITY itself (20).

As such while changes within the islet microvasculature during pre-symptomatic T1D still requires characterization, there is feasibility to translate the methods here to human T1D.

### Summary

In summary, we demonstrate that highly novel nanodroplet ultrasound contrast agents specifically accumulate within the islets of animal models of T1D, where this accumulation can be measured via contrast-enhance ultrasound following nanodroplet vaporization. This accumulation can inform both on the level of insulitis across the pancreas and the rapid progression to T1D. This method may be translatable to human pre-symptomatic T1D to guide therapeutic intervention to prevent T1D.

## Materials and Methods

### Animals

All animal procedures were performed in accordance with guidelines established by the Institutional Animal Care and Use Committee of the University of Colorado Anschutz Medical campus. Female NOD mice were purchased from Jackson Laboratories (Bar Harbor, ME) at age 8 weeks, and imaged at 10 weeks of age. Female Rag1ko animals were bred inhouse, and imaged at 10 weeks of age. Throughout the study, animals were monitored weekly for blood glucose concentration utilizing a blood glucometer (Bayer).

### Nanodroplet synthesis

Microbubbles were formed using previously published methods (29). Briefly DPPC and DSPE-PEG2000 (5:1 weight ratio) were dissolved in chloroform and evaporated into a lipid film. This film was reconstituted into PBS, the mixture sonicated with an ultrasonic probe at low power while bubbled with perfluorobutane, to generate a suspension of gas-filled microbubbles. The solution was then placed in an ice bath until the solution temperature was 23°C. 4-5 μm size isolated bubbles were then formed via a series of differential centrifugation steps and PBS washing. Size distributions and concentrations of the microbubbles were measured by laser light scattering and obscuration using the Accusizer 780A.

Condensation of microbubbles into nanodroplets (19, 30) was achieved by cooling a suspension of microbubbles in PBS down to ~1°C in an isoflurane bath. A high pressure (~10 bar) of perfluorobutane was then applied until the solution turned transparent. Size distributions and concentrations of the nanodroplets were measured using the Accusizer 780A.

Polydisperse nanodroplets were formed initially by reconstituting a DPPC:DSPE-PEG2000 film in PBS as above. This solution was then mechanically agitated using a Vialmix. Fluorescent labelling of nanodroplets by DiO was achieved by diluting the lipid film mixture with 6μm DiO.

### Contrast-enhanced ultrasound (CEUS) imaging

General anesthesia was established with isoflurane inhalation for ~40-50 minutes for all mice imaged. A custom made 27G ½” winged infusion set (Terumo BCT, Lakewood, CO) was attached to a section of polyethylene tubing (0.61 OD x 0.28 ID, Warner Instruments) and inserted into a lateral tail vein. Abdominal fur was removed using depilatory cream and ultrasound gel was placed to match the acoustic impedance between the mouse and transducer. Electrocardiogram, respiration rate and body temperature were monitored by foot pad electrodes on the ultrasound platform. Body temperature and respiration rate were monitored throughout the imaging session.

A VEVO 2100 small animal high-frequency ultrasound machine (Visual Sonics, Fujifilm, Toronto, Canada) was used for the experiments. A MS250 linear array transducer at 18 MHz was used. B-mode imaging was performed prior to ND infusion to identify the pancreas body in relation to the stomach, kidney, and spleen (Fig. 1b). Following successful identification, a region of interest was defined around individual organs (pancreas or kidney) and sub-harmonic contrast mode was initiated. Acquisition settings were set to: transmit power 10% (MI=0.12); frequency 18 MHz; standard beamwidth; contrast gain 30 dB; 2D gain 18 dB; acquisition rate 26 frames/second. Gating to remove breathing movement was carried out manually or using custom scripts MATLAB (Mathworks, Natick, MA).

Background images were acquired for 3 minutes, at 30 second intervals, prior to nanodroplet infusion. Nanodroplets (ND) were then infused as a single bolus of 100 μl solution 2×10^8^ ND/ml into the lateral tail vein. Following a 25-minute pause images were acquired for 10 minutes at 30 second intervals.

The vaporized nanodroplet contrast signal was background subtracted by the mean contrast intensity taken prior to nanodroplet infusion. Each time course was normalized to the pre-infusion background values to provide the S-B/B. B-mode imaging was used to identify the pancreas and kidney and regions of interest were placed on the organs. The regions of interest were manually adjusted prior to analysis to avoid small regions of high contrast saturation in the background images.

### Histology and Insulitis Morphology

For assessment of insulitis, animals were anesthetized by I.P. injection of Ketamine (80 mg/kg) and Xylazine (16 mg/kg) until no longer reactive to toe pinch. The pancreata were isolated and mice were euthanized by exsanguination and/or Bilateral thoracotomy. Pancreata were fixed in paraformaldehyde at 4°C in a mechanical rocker overnight and embedded in OCT blocks the following day. 8 μm sections were taken at three tissue depths. Alternating slices were stained with hematoxylin and eosin (H&E) for evaluation of islet T cell infiltration. Images were acquired on an Eclipse-Ti wide field microscope with a 20X 0.75 NA Plan Apo objective with a color CCD camera.

Images of islets were scored based on the extent of infiltration/insulitis: grade 0, no insulitis; grade 1, peri-insulitis with immune infiltrate bordering; grade 2, immune infiltrate penetrating the islet, covering <50% of the islet area; grade 3, immune infiltrate penetrating the islet, covering >50% of the islet area. A minimum of three different tissue depths and at least 30 non-overlapping islets per animal were analyzed. Weighted averages were calculated for each animal.

For assessment of nanodroplet coverage 10w female NOD mice (JAX) received a single bolus injection of 100 μl solution of 2×10^8^ ND/ml of DiO-labeled nanodroplets. Following contrast imaging, mice were anesthetized by I.P. injection of ketamine (80 mg/kg) and xylazine (16 mg/kg). The pancreas was dissected and fixed in 4% PFA on ice for 1 hour and cryoprotected in 30% sucrose overnight or until the tissue sank. Pancreata were embedded in OCT medium, frozen in cryomolds, and cryo-sectioned at 8μm sections, as above. Sections were imaged on an LSM800 confocal microscope (Zeiss), with at 488nm excitation using a x20 0.8 NA objective and pinhole settings to provide 1μm z-section thickness throughout the tissue depth. Separate images were taken of exocrine tissue at locations anatomically isolated from the islets. DiO coverage was calculated in MATLAB as the area of DiO positive pixels (pixels with fluorescence intensity significantly above the background fluorescence intensity) across the islet (or other defined region) and expressed as a fraction of total islet area.

## Acknowledgments

Richard KP Benninger (University of Colorado) is the guarantor of this work and, as such, had full access to all the data in the study and takes responsibility for the integrity of the data and the accuracy of the data analysis. All authors acknowledge that no conflict of interest exists. This work was supported by Juvenile Diabetes Research Foundation Grants 1-INO-2017-435-A-N, 5-CDA-2014-198-A-N; and NIH grants R01 DK102950, R01 DK106412 (to RKPB); by NIH grant F31 DK121488 (to DGR); by NIH grant TL1 TR002533 (MC); by NIH grant R01CA195051 (MAB). The funders had no role in the study design, data collection and analysis, decisions to publish, or preparation of the manuscript.

## Author Contributions

DR designed and performed experiments, analyzed data, wrote the manuscript; VP performed experiments and analyzed data; AU performed experiments; MC performed experiments; MAB conceived of the idea, designed experiments and edited the manuscript; RKPB conceived of the idea, designed experiments, performed experiments and wrote the manuscript.

## Supplemental Materials

**Supplemental Figure S1.**
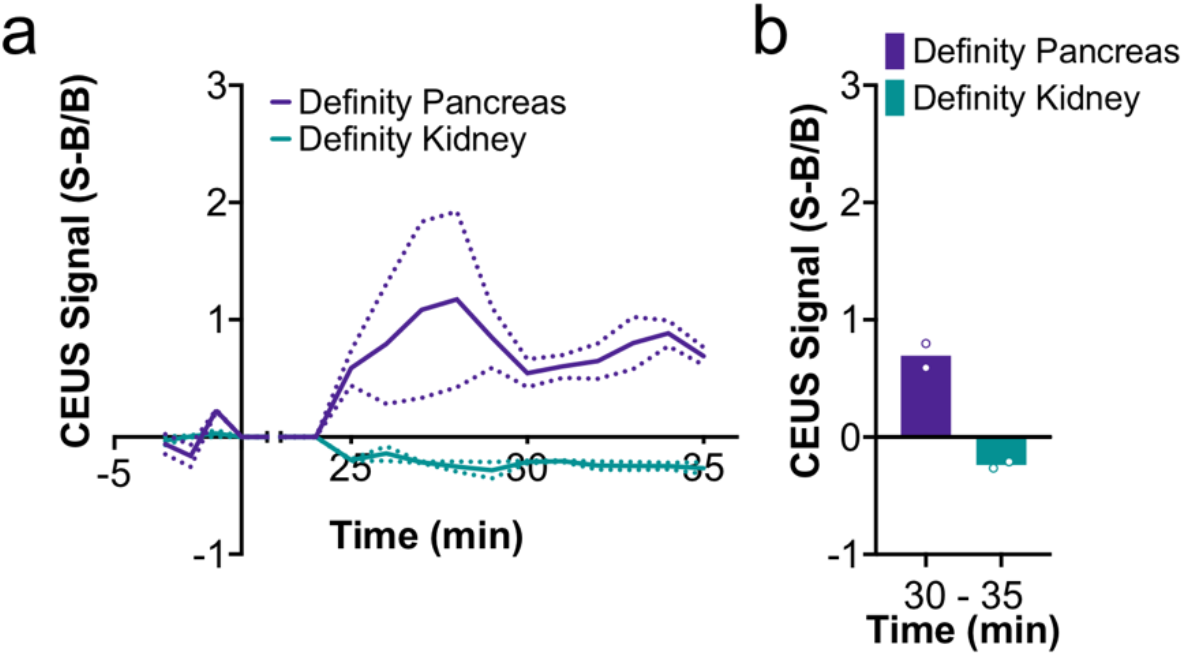
Polydisperse nanodroplets show increased contrast within the pancreas prior to T1D. (**A**) Mean time-course of contrast signal in the pancreas of 10 week old female NOD mice with NDs made from poly-dispersed microbubbles. (B) Mean contrast signal averaged between 30-35 minutes following ND infusion with NDs made from poly-dispersed microbubbles. Dashed lines in A represent 95% CI.

**Supplemental Figure S2.**
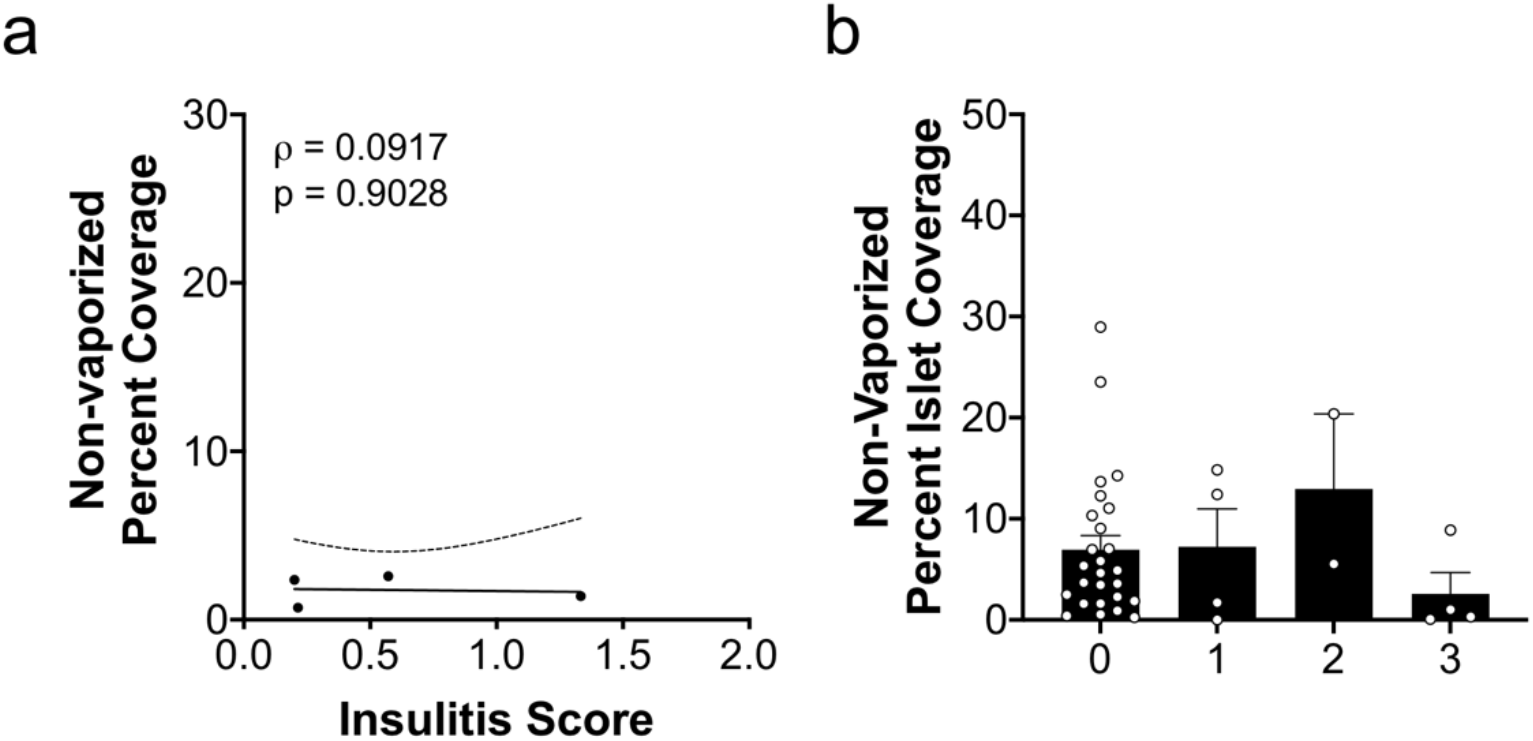
Histological assessment of non-vaporized DiO labelled nanodroplets in the islets in T1D. (A) Scatterplot of nanodroplet-DiO fluorescent coverage of islets within the pancreas against the mean insulitis score within the pancreas, for NOD mice without ultrasonic vaporization. (B) Mean nanodroplet-DiO fluorescent coverage of non-vaporized islets within the pancreas that show insulitis scores of 0, 1, 2, or 3. Error bars in B represent s.e.m. Trend line in A indicates linear regression with 95% confidence intervals. ANOVA was performed in B. A mixed-effects model was used to assess the statistical significance and generate the regression in A. Data in B represents 36 islets from 4 NOD mice.

